# High throughput characterization of genetic effects on DNA:protein binding and gene transcription

**DOI:** 10.1101/270991

**Authors:** Cynthia A. Kalita, Christopher D. Brown, Andrew Freiman, Jenna Isherwood, Xiaoquan Wen, Roger Pique-Regi, Francesca Luca

## Abstract

Many variants associated with complex traits are in non-coding regions, and contribute to phenotypes by disrupting regulatory sequences. To characterize these variants, we developed a streamlined protocol for a high-throughput reporter assay, BiT-STARR-seq (Biallelic Targeted STARR-seq), that identifies allele-specific expression (ASE) while accounting for PCR duplicates through unique molecular identifiers. We tested 75,501 oligos (43,500 SNPs) and identified 2,720 SNPs with significant ASE (FDR 10%). To validate disruption of binding as one of the mechanisms underlying ASE, we developed a new high throughput allele specific binding assay for NFKB-p50. We identified 2,951 SNPs with allele-specific binding (ASB) (FDR 10%); 173 of these SNPs also had ASE (OR=1.97, p-value=0.0006). Of variants associated with complex traits, 1,531 resulted in ASE and 1,662 showed ASB. For example, we characterized that the Crohn’s disease risk variant for rs3810936 increases NFKB binding and results in altered gene expression.

---

Genome wide association studies (GWAS) have identified thousands of common genetic variants associated with complex traits, including normal traits and common diseases. Many GWAS hits are in non-coding regions, so the underlying mechanism leading to specific phenotypes is likely through disruption of gene regulatory sequence. Quantitative trait loci (QTLs) for molecular and cellular phenotypes [1], such as gene expression (eQTL) [2, 3, 4, 5, 6], transcription factor binding [7], and DNaseI sensitivity (dsQTL) [8] have been crucial in providing strong evidence and a better understanding of how genetic variants in regulatory sequences can affect gene expression levels [9, 6, 10, 11]. In recent work, we were able to validate 48% of computationally predicted allelic effects on transcription factor binding through traditional reporter assays [12]. However, traditional reporter assays are limited by the time and the cost of testing variants one at a time.

Massively parallel reporter assays (MPRA) have been developed for the simultaneous measurement of the regulatory function of thousands of constructs at once. For MPRA, a pool of synthesized DNA oligos containing a barcode at the 3’UTR of a reporter plasmid is transfected into cells, and transcripts are isolated for RNA-seq. The number of barcode reads in the RNA over the number of barcode reads from the plasmid DNA is used as a quantitative measure of expression driven by the synthesized enhancer region [13, 14, 15, 16, 17]. An alternative to MPRA is STARR-seq (self-transcribing active regulatory region sequencing) [18], whose methods involve fragmenting the genome and cloning the fragments 3’of the reporter gene. The approach is based on the concept that enhancers can function independently of their relative positions, so putative enhancers are placed downstream of a minimal promoter. Active enhancers transcribe themselves, with their strength quantified as the amount of RNA transcripts within the cell. Because they do not use separate barcodes, STARR-seq approaches have streamlined protocols that allow for higher throughput.

Recently, high-throughput assays have been used to assess the enhancer function of genomic regions [18, 19], the allelic effects on gene expression for naturally occurring variation in 104 regulatory regions [20], fine-map variants associated with gene expression in lymphoblastoid cell lines (LCLs) and HepG2 [21], and fine-map variants associated with red blood cell traits in GWAS [22]. In addition to using reporter assays to measure enhancer function on gene expression, there are several methods to directly measure binding affinity of DNA sequences for specific transcription factors. These methods include Spec-seq [23], EMSA-seq (electrophoretic mobility shift assay-sequencing) [24], and BUNDLE-seq (Binding to Designed Library, Extracting, and sequencing) [25]. In these assays, synthesized regions are combined *in vitro* with a purified transcription factor. The bound DNA-factor complexes are then isolated by polyacrylamide gel electrophoresis (PAGE). Extracting the DNA from upper (bound complex) and lower (unbound DNA) bands and sequencing of the derived libraries allows for quantification of the binding strength of regulatory regions. While BUNDLE-seq compared binding and reporter gene expression, and EMSA has been previously used to ascertain allelic effects, none of the high-throughput EMSA methods have been previously used to determine allelic effects on binding.

We have developed a method called BiT-STARR-seq (Biallelic Targeted STARR-seq) to test for allele specific effects in regulatory regions (Figure 1a). BiT-STARR-seq applies the streamlined protocol of STARR-seq to thousands of synthesized oligos targeting independent genomic regions, to create the simplest experimental protocol for high-throughput reporter assays to date. The method also includes the incorporation of unique molecular identifiers (UMIs) during cDNA synthesis, that allows for the removal of duplicates created during library preparation. We used BiT-STARR-seq to test 43,500 regulatory variants, including variants predicted to disrupt transcription factor binding (CentiSNPs [12]) for 874 transcription factors, as well as other regulatory variants [5, 4, 26, 12]. We then adapted BUNDLE-seq to analyze allele-specific binding (ASB) for NFKB (p50) and validate the molecular mechanism underlying the allele-specific effects measured in the BiT-STARR-seq assay. We denote this new method BiT-BUNDLE-seq (Biallelic Targeted BUNDLE-seq). To the best of our knowledge, this is the first use of any high-throughput EMSA to consider allele-specific binding for regulatory regions. Our results demonstrate that high-throughput EMSA approaches complement allele-specific analyses in MPRA/STARR-seq assays, thus providing an effective strategy to dissect the molecular mechanism linking regulatory variants effects on binding and on expression. Our method is especially well suited to test in parallel thousands of computationally prioritized variants that are also associated with complex traits.

We selected different categories of regulatory variants for this study including eQTLs [5, 4], CentiSNPs [12], ASB SNPs [12], variants associated with complex traits in GWAS [26], and negative ASB controls [12] for a total of 50,609 SNPs. We designed two oligos targeting each of the alleles for a SNP, with inserts 230bp long synthesized by Agilent to contain the regulatory region and the SNP within the first 150bp. We also included the use of unique molecular identifiers (UMIs), added during cDNA synthesis. With these random UMIs we are in effect tagging identifiable replicates of the self-transcribing construct, which improves the analysis of the data by accounting for PCR duplicates. Our protocol also has the advantage of being highly streamlined. Unlike STARR-seq, our method does not require preparation of DNA regions for use in the assay, such as whole genome fragmentation [18], or targeting regions [19, 27], while, similar to STARR-seq, it requires only a single cloning and transformation step. Because the UMIs are inserted after transfection, there are no additional bottleneck issues (due to library complexity) in the cloning and transformation steps.

**Figure 1:**
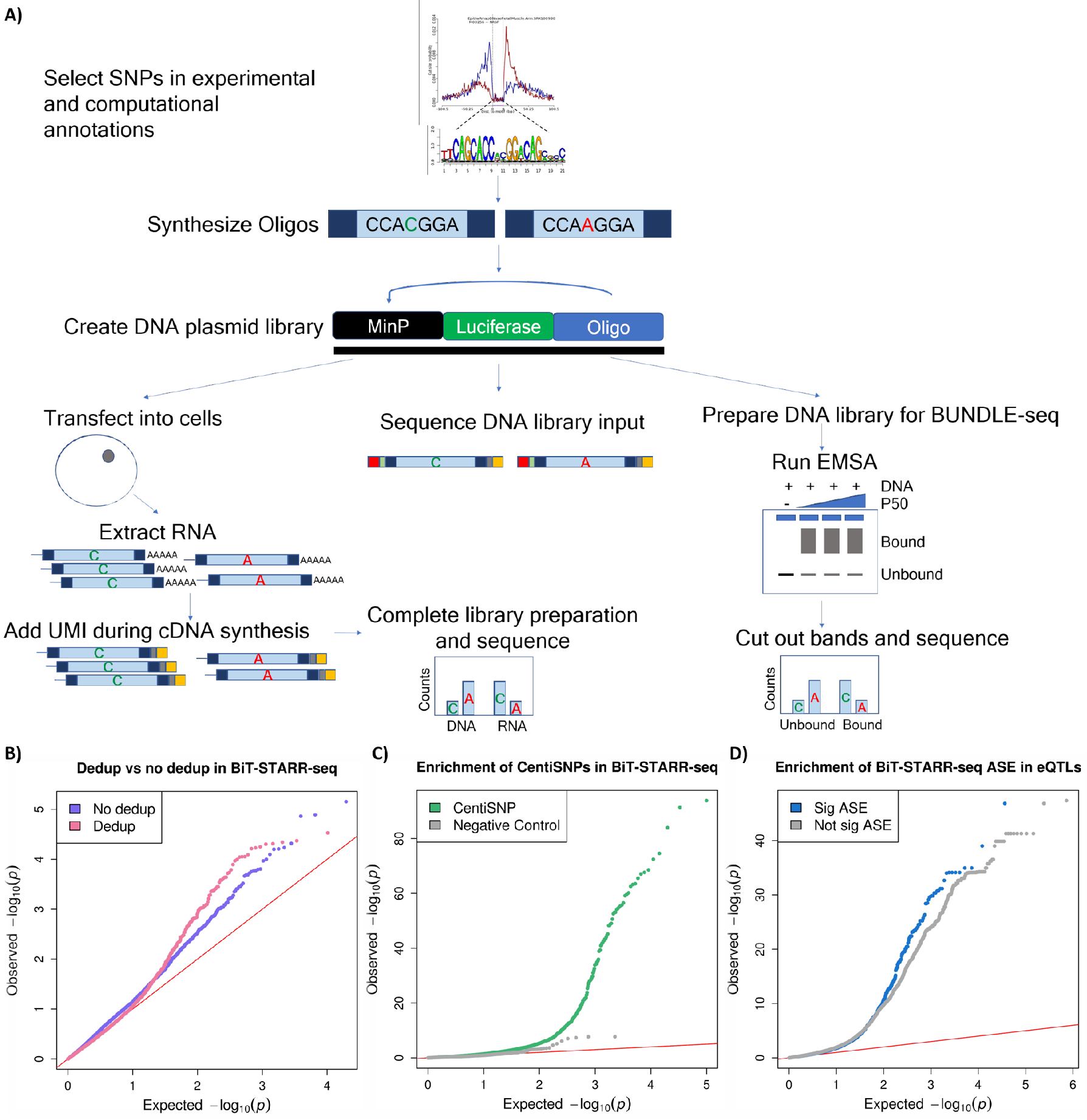
BiT-STARR-seq and BiT-BUNDLE-seq identify regulatory variants in non-coding regions. A) Experimental outline. Oligos targeting the regulatory regions of interest (and either reference or alternate alleles) are designed to contain, on their ends, 15bp matching the sequencing primers used for Illumina NGS. The DNA library is used both in the BiT-STARR-seq and BiT-BUNDLE-seq experiments. UMIs are added during cDNA synthesis for the BiT-STARR-seq RNA-seq library and prior to PAGE in the BiT-BUNDLE-seq protocol. B) QQplot depicting the *p*-value distributions from QuASAR-MPRA for a single experimental replicate processed without removing duplicates (purple) or after removing duplicates using the UMIs (pink). C) QQplot depicting the *p*-value distributions from the ASE test performed using QuASAR-MPRA on all replicates after removing duplicates. CentiSNPs are in (green)[12] while SNPs in the negative control group are in (grey). D) QQplot depicting the *p*-value distributions for eQTLs from [28]. SNPs with significant ASE (FDR 10%) are in (blue) or not significant ASE are in (grey).

We generated 7 replicates of the DNA library, which were highly and significantly correlated (Figure S1 Spearman′s *ρ* = (0.97, 0.98), *p*-value<0.01). The DNA library was then transfected in LCLs (9 biological replicates) and we were able to recover a total of 43,500 testable SNPs (see methods for RNA counts filter). Read counts for the 9 biological replicates were highly correlated (Figure S2 Spearman’s *ρ* = (0.35, 0.72), *p*-value<0.01). To identify SNPs with allele-specific effects, we applied QuASAR-MPRA [29]. For each SNP, the reference and alternate allele counts were compared to the DNA proportion in the plasmid library. We identified a total of 2,720 SNPs with ASE from the combined replicates (FDR 10%) (Table S6). To investigate the importance of UMIs in this experimental approach, we re-analyzed our data without removing duplicates. For the combined replicates, inflation is greatly increased (from 1.10 to 1.73). Using UMIs in STARR-seq approaches is particularly relevant when a small number of replicates is used. For example, in the extreme case when a single replicate is considered, inflation increases slightly (from 0.92 to 1.18) and the number of detected ASE is about 4x reduced (Figure 1b).

SNPs with ASE are significantly enriched for variants predicted to impact transcription factor binding (CentiSNPs) [12] (Figure 1c, Table S1a, Fisher′s exact test *p*-value=4.55 × 10^−06^, OR=2.49). Additionally, SNPs with ASE are enriched for low p-values in an eQTL mapping study performed in immune cells [28] (Figure 1d, Table S1c), thus confirming that our synthetic oligos can reproduce allele-specific regulatory effects observed in a native (non-episomal) cellular context.

The CentiSNP annotation is informative of the specific transcription factor motif being disrupted by a SNP. By leveraging this information, we were able to analyze allelic effects for specific transcription factor motifs (Table S7) (Figure 2a). Additionally, by combining the ASE results with the direction of the motif, we can characterize whether the motif is active in both directions or only in one direction. In other words if for a factor ASE tends to occur only in one direction, this would suggest a directional effect on the regulatory function. We found that when both alleles are covered in both directions, the allelic effects on gene expression tend to agree in direction and magnitude. If we categorize these directional allelic effects per motif, we do not observe major differences with the notable exception of CTCF (Figure 2b). Specifically, we find that SNPs in footprints for CTCF are significantly enriched (Fisher′s exact test OR=1.57, Bonferroni p=0.02) for ASE when the direction of transcription of the reporter gene is opposite to that of the motif strand (Figure 2c).

**Figure 2:**
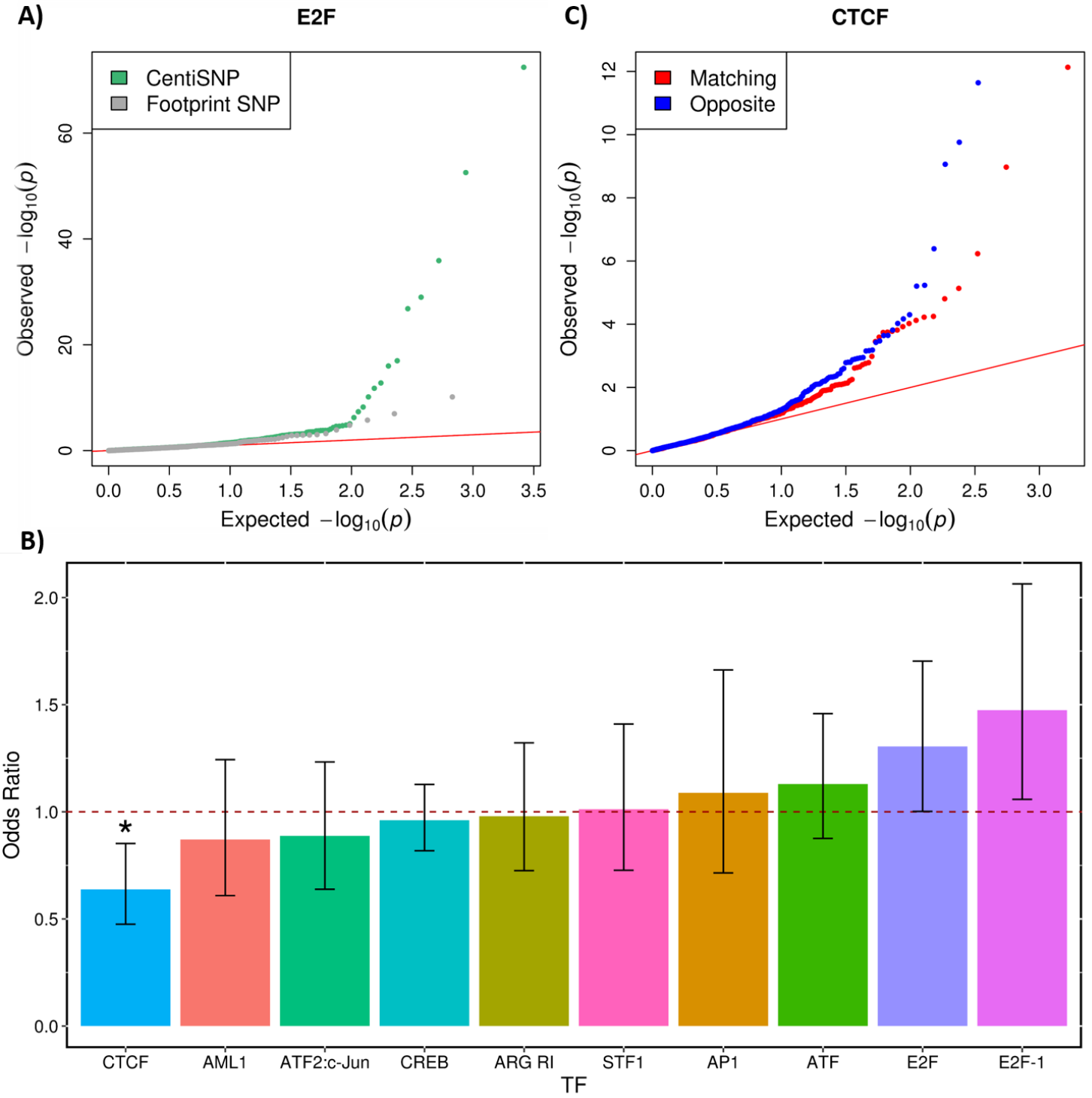
ASE for individual transcription factors. A) QQplot depicting the ASE *p*-value distributions from QuASAR-MPRA, for SNPs overlapping with E2F footprint annotations. SNPs predicted to alter binding (CentiSNPs) are represented in green, while SNPs that are in E2F but predicted to have no effect on binding are in grey. B) Enrichment for ASE in individual transcription factor binding sites calculated when motif strand matched the BiT-STARR-seq oligo transcription direction. Odds ratio (y axis) for each transcription factor tested (x axis) is shown in the barplot, error bars are the 95% CI from the Fisher’s exact test. Odds ratios below the dotted line represent enrichment for opposite direction oligo/motif configuration. Stars are shown above significant results (Bonferroni adjusted *p*-value <0.05). C) QQplot depicting the ASE *p*-value distributions from QuASAR-MPRA, overlapping with CTCF footprint annotations. In red are the SNPs where the motif strand matches the BiT-STARR-seq oligo direction relative to the TSS, in blue where the motif strand is the opposite of the BiT-STARR-seq direction.

In order to understand the effect of a regulatory variant on complex traits it is necessary to first dissect the molecular function that is impacted by the nucleotide change. The CentiSNP prediction provides specific hypotheses for allelic effects on transcription factor binding that can be directly tested experimentally. Further matching ASB to ASE identified in BiT-STARR-seq would provide a complete molecular mechanism, from predicted binding effects, to experimentally validated binding effects, to validated effects on expression. Due to the enrichment of CentiSNPs among SNPs with ASE in BiT-STARR-seq, we performed BiT-BUNDLE-seq to validate their effect on transcription factor binding. This is a new and efficient extension of high throughput reporter assays, since it uses the same input DNA library. We performed BiT-BUNDLE-seq with purified NFKB-p50 (at three different concentrations), which is an important regulator of the immune response in LCLs and other immune cells [30, 31, 32]. Previous studies have successfully identified ASB from ChIP-seq for NFKB in LCLs [33, 34, 35, 7, 36, 37] and NFKB footprints are induced in response to infection [38]. Additionally, NFKB was found to be 50 fold enriched for reQTLs from response to Listeria and Salmonella [28].

We first analyzed NFKB-p50 binding between the bound and unbound libraries and identified 9,361 significantly (logFC>1 and FDR<1%) over-represented regions in the bound library (Table S3). As expected, these regions were enriched (OR=11.70,13.75, *p*-value=7.95 × 10^−27^,1.23 × 10^−15^) for NFKB footprints (Figure 3a, S3), with a higher portion of these regions in the mid concentration of NFKB-p50 as compared to the low or high concentrations (Figure 3b). This is probably because the low concentration doesn′t capture all of the NFKB-p50 binding, whereas the high concentration likely results in non-specific binding. We then used ΔAST [39] to identify ASB in the bound library (as compared to the unbound library), and combined the replicates using Stouffer′s method (See Methods). We successfully identified 386 SNPs at low concentration, 797 SNPs at mid concentration, and 894 SNPs at high concentration with significant ASB at 10% FDR (Figure 3d), for a total of 2,951 SNPs when aggregating all experiments (Table S1b). When we considered these ASB SNPs in combination with the ASE results from the BiT-STARR-seq, we found that SNPs with ASE are enriched for NFKB-p50 ASB (Figure 3e, Fisher′s exact test (OR=2.04, *p*-value=1.51 × 10^−16^)). This confirms our hypothesis that disruption of NFKB-p50 binding is one of the mechanisms underlying allele-specific expression in our dataset.

**Figure 3:**
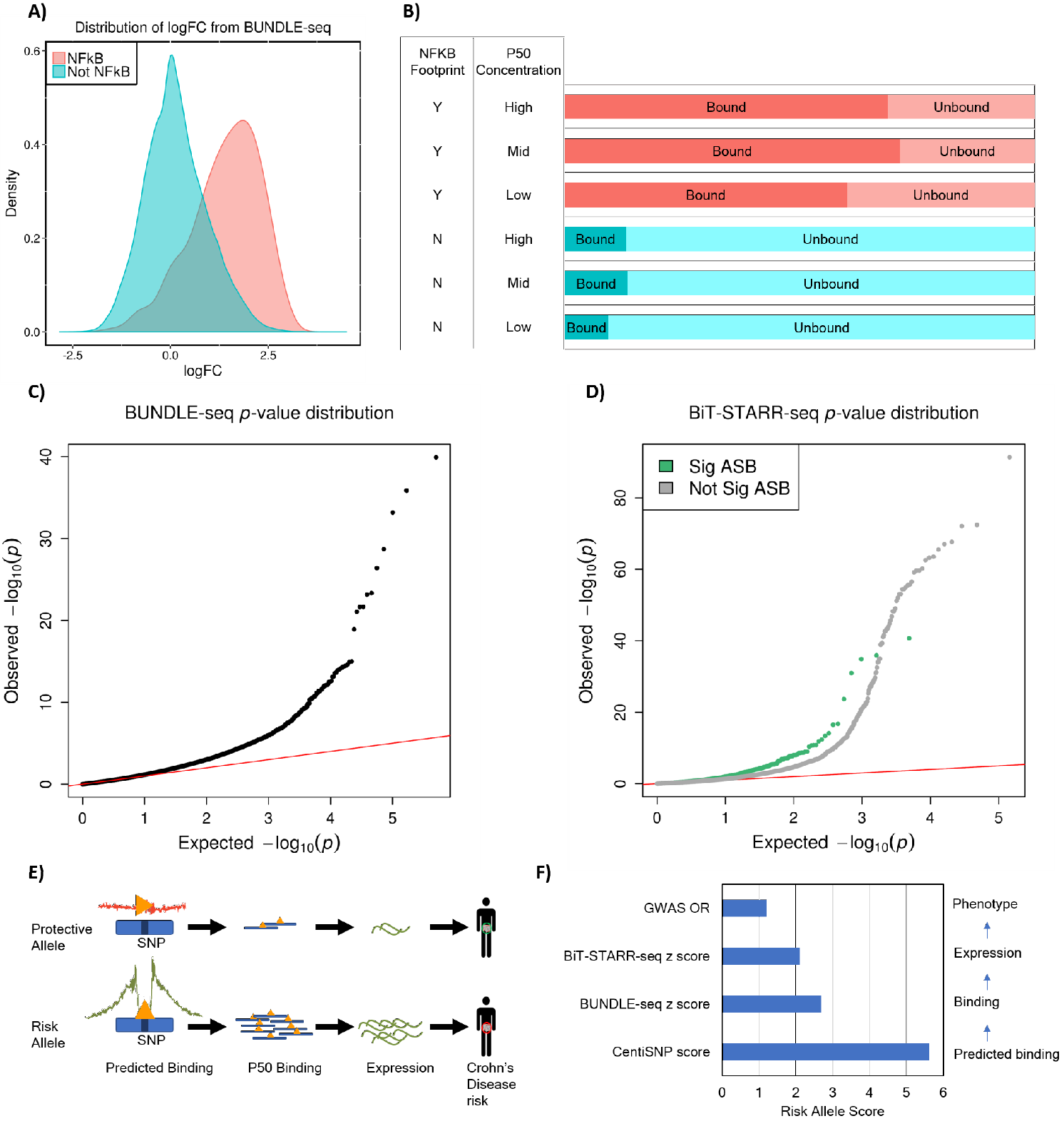
Allele-specific binding for NFKB-p50. A) Density plot of the logFC (from DESeq2) between bound and unbound DNA fractions from the BiT-BUNDLE-seq experiment. In red are the regions containing a SNP in a NFKB footprint, in blue the regions containing a SNP in footprints for other transcription factors. B) Barplot representing the number of independent enhancer regions in bound (dark color, DESeq2 logFC>1 and FDR<1%) and unbound (light color) DNA. NFKB-p50 concentration and presence of a NFKB footprint are indicated in the two columns on the left of the panel. C) QQplot depicting the *p*-value distributions from testing for ASB signal specific to the bound DNA fraction using ΔAST. D) QQplot depicting the ASE *p*-value distribution from QuASAR-MPRA for SNPs with significant (FDR<10%) (green) or not significant (grey) ASB in the BiT-BUNDLE-seq experiment. E) Integration of prediction, BiT-BUNDLE-seq, BiT-STARR-seq and GWAS results for Crohn’s disease risk variant rs3810936. Triangles represent transcription factors. F) Comparing allelic effects from computational prediction to phenotype for rs3810936. Predicted log odds score is the reference prior log odds - alternate log odds from the CentiSNP annotation. BiT-BUNDLE-seq z score is the z score from the meta-analysis of ASB from all 3 concentrations of NFKB-p50. BiT-STARR-seq z score is the z score from meta-analysis of ASE for nine experimental replicates. GWAS OR is the odds ratio from rs3810936 alternate allele with Crohns disease [40, 41, 42, 43]. All scores are signed relative to the risk allele, which is the alternate allele.

We used ASB and ASE in combination with transcription factor binding motifs to assign mechanistic function to putatively causal SNPs linked to complex traits. We found 2,054 CentiSNPs with ASB (*p*-value<0.05) and 1,769 CentiSNPs with ASE (*p*-value<0.05) associated to a complex trait in the GWAS catalog (Table S5,S6) or with a PPA>0.1 from fgwas [12]. We were able to dissect the molecular mechanism for rs3810936, a variant associated with risk for Crohn′s disease in multiple populations [40, 41, 42, 43] (Figure 3e,f). This variant is a CentiSNP for the factor Hmx3 (Nkx5-1) and we find ASB for NFKB (*p*-value=0.006) in the BiT-BUNDLE-seq assay and ASE (*p*-value=0.034) in both directions in the BiT-STARR-seq. This SNP is a synonymous variant in gene *TNFSF15* (also known as *TL1A*), which encodes for a cytokine that belongs to the tumor necrosis factor (TNF) ligand family. Increased TL1A expression has been reported in inflamed Crohn′s disease tissue compared with non-inflamed areas, and in ulcerative colitis patient serum [44, 45, 46]. TL1A gives costimulatory signals to activated lymphocytes through binding to death-domain receptor 3 (DR3) [47] which induces the secretion of IFN-gamma [48, 49]. This gene modulates Th-1 and Th-17 [45, 50], creating an immunological state that leads to the mucosal inflammation of Crohn′s disease. Stimulation of the TL1A pathway, in monocytes and T cells from patients carrying the disease-associated *TL1A* SNPs, showed higher levels of TL1A expression, therefore aberrant TL1A expression may be a factor driving IBD development [51, 52]. This gene additionally has been found to be downregulated in response to dexamethasone [39], a corticosteroid used to treat many inflammatory and autoimmune conditions. While this variant is not found in ChIP-seq from ENCODE, ENCODE studies used RELA (p65) for NFKB, where our study used p50. We therefore identify a novel variant that disrupts binding of NFKB-p50, where the alternate allele (C) has increased binding. This leads to a increase in gene expression for the alternate allele, which is also the risk allele for Crohn′s disease (OR=1.21, *p*-value=1 × 10^−15^).

The recent adaptation of MPRA to investigate ASE allows for validation of regulatory variants in transcription factor binding sites, which have been shown to be functionally relevant to fine map eQTLs and GWAS signals. However, the use of functional genomics to select relevant regions prior to experimental validation can reduce the number of sites it is necessary to validate. We developed a high throughput reporter assay that synthesizes these selected regions, clones them in 3’of the reporter gene, and includes the addition of a UMI during cDNA synthesis. This is the most streamlined protocol to date, and allows for removal of PCR duplicates which reduces noise in the data for greater power to detect ASE.

Our results show that using existing annotations to prioritize regulatory variants for high-throughput reporter assays is an effective strategy. The CentiSNP annotation in particular contains information that can be used to analyze ASB/ASE for individual transcription factor motifs and investigate potential molecular mechanisms of action. We found that direction is an important factor in the case of CTCF, most likely due to how CTCF functions as an insulator between the enhancer and the promoter when they are in anti-parallel directions. While enhancers can exert their regulatory function in either direction [53], insulators are only active if they can bind the DNA in specific arrangements relative to the promoter [54, 55]. Previous studies have shown that CTCF, a well characterized insulator, has binding sites at the anchors of chromatin loops.

These are arranged in forward-reverse orientations [53, 54, 55, 56, 58], where the relative positions and orientations of the binding sites are important for the mechanism of action [53]. In our case, the interaction could be mediated either by the basal transcriptional machinery at the TSS or also an additional weak CTCF binding site (M01259) that is present in the promoter and could help to establish a DNA loop. However, there may be alternative explanations for this result as reporter assays may not reflect the native regulatory landscape in human cells [59, 60]. Generally, caution should be used in interpreting reporter assay gene expression differences across cell types, because transfection may perturb the cell state. However, it is important to highlight that any trans-acting effects (e.g., promoter strength, IFN-1 response activation) should affect both alleles similarly and therefore should not induce false positives in the allele-specific signal.

We used our library of oligos also in a BiT-BUNDLE-seq assay for identification of ASB for NFKB-p50. This is a novel approach to combine ASB and ASE identification in high throughput assays using the same sequences. Our results show that this integration is a useful approach to validate the molecular mechanism for specific transcription factors.

Allelic effects on transcription factor binding and gene expression are not always concordant. This is the case, for example, of an allele that increases binding of a factor with repressing activity on gene expression. For example, we identified regulatory variants where there is increased binding for NFKB-p50, but decreased expression. These variants are in regions enriched for the CREB motif, and CREB has been shown to antagonize NFKB binding [61, 62]. These regulatory events are likely to be captured in the BiT-STARR-seq assay, which is performed in LCLs where both CREB and NFKB are active. These results highlight that multiple type of assays are necessary to capture the detailed molecular mechanism of gene regulation. Additionally, integration with GWAS can identify and further characterize the molecular mechanisms linking causal genetic variants with complex traits.

## Methods

### Oligo selection and design

Table S1 reports the annotations we have considered with their sources. These included: SNPs predicted to alter transcription factor binding in LCLs and HepG2 (CentiSNPs, [63]), LCL eQTLs fine-mapped in [5], liver eQTLs [4], significant fgwas SNPs in transcription factor binding motifs for 18 complex traits [12], significant fgwas S NPs for base models of functional annotations for 18 complex traits [26], ASB SNPs, and strong enhancers with no predicted ASB [63]. CentiSNP is an annotation that we recently developed [12], and that uses the CENTIPEDE framework [64] to integrate DNase-seq footprints with a recalibrated position weight matrix (PWM) model for the sequence to predict the functional impact of SNPs in footprints. SNPs in footprints “footprint-SNPs” are further categorized using CENTIPEDE hierarchical prior for each allele as “CentiSNP” if the prior relative odds for binding are >20. Fasta sequences with a window of 99 (on each side of the SNP) on the bed file were grabbed using seqBedFor2bit, and 15bp matching sequencing primers used for Illumina NGS were added to each end. Each regulatory region was designed to have two oligos: one for each of the alleles. A second list of the fasta sequences without the primer ends was generated to use as a custom reference genome, then converted to fastq using faToFastq. The full SNP list was aligned to the hg19 genome with BWA mem [65], removing the regions with a quality score less than 20. The full SNP list was also aligned to the custom reference genome, and then filtered for a quality score of 190. A total of 39,366 indexes were randomly generated to match this pattern: RDHBVDHBVD. This sequence was chosen to limit the longest possible polyACGT run at any position to 3 nucleotides, and avoid a G in the first and last position (corresponding to a dark cycle on the Illumina NextSeq500).

### Oligo synthesis and amplification

DNA inserts 230bp long, corresponding to 200bp of regulatory sequence, were synthesized by Agilent to contain the regulatory region and the SNP of interest within the first 150bp. We performed a first round of PCR using Phusion High-Fidelity PCR Master Mix with HF Buffer (NEB) and primers [F transposase and R transposase] with cycling conditions: 98°C for 30s, followed by 4 cycles of 98°C for 10s, 50°C for 30s, 72°C for 60s, followed by 6 cycles of 98°C for 10s, 65°C for 30s, 72°C for 60s, followed by 72°C for 5 min. This reaction was used to double strand the oligos and complete the sequencing primers. The PCR product was run on a 2% agarose gel, extracted and purified with the NucleoSpin Gel and PCR Clean-Up Kit (Clontech). A subsequent round of PCR amplified the material using the same reaction as in the first round of PCR, but with cycling conditions: 98°C for 30s, followed by 15 cycles of 98°C for 10s, 65°C for 30s, 72°C for 60s, followed by 72°C for 5min. The PCR product was purified as described above.

### Cloning Regulatory regions into pGL4.23

A recent study demonstrated that the ORI can be an active promoter in pGL4.23 plasmids and can function as a stronger promoter in the absence of other promoter sequences [59]. Here we used a design that includes a minimal promoter, thus potentially missing some signal from the weakest enhancer sequences in our library. However, as we focus on allele-specific enhancer activity, the presence of a minimal promoter in addition to the ORI, should affect both alleles similarly and should not induce false positives in the allele-specific signal. Plasmid pGL4.23 (Promega) was linearized using CloneAmp HiFi PCR Premix (Clontech), primers [STARR F SH and STARR R SH], and 35 cycles of 98°C for 10s, 60°C for 15s, and 72°C for 5s. The PCR product was purified on a 1% agarose gel as described above. Inserts were cloned into the linear plasmid using standard Infusion (Clontech) cloning protocol. Clones were transformed into XL10-Gold Ultracompetent Cells (Agilent) in a total of 7 reactions. These reactions were pooled and grown overnight in 500ml LB at 37°C in a shaking incubator. DNA was extracted using Endofree maxiprep kit (QIAgen).

### Transfection of library

Previous studies [59, 60] have found that transfection, especially from nucleofection, can lead to activation of IFN-1 response, which may complicate comparison of enhancer activities between different cell types. In our study design, allelic effects are measured and contrasted within the same cell type, thus any trans-effect is inherently controlled. Furthermore, in LCLs the immune response is already activated because of EBV transformation. DNA library was transfected into LCLs using standard nucleofection protocol, program DS150, 3*μ*g of DNA and 7.5 × 10^6^ cells. A total of 3 sets of transfections were done in triplicate cuvettes, then pooled. We performed nine biological replicates of the transfection from 7 independent cell growth cultures. After transfection, cells were incubated at 37°C and 5% CO2 in RPMI1640 with 15%FBS and 1% Gentamycin for 24h. Cell pellets were then lysed using RLT lysis buffer (QIAgen), and cryopreserved at −80°C.

### Library preparation

RNA-libraries. Thawed lysates were split in three aliquotes and total RNA was isolated using RNeasy Plus Mini Kit (QIAgen). Poly-Adenylated RNA was selected using Dynabeads mRNA Direct Kit (Ambion) using the protocol for total RNA input. RNA was reverse transcribed to cDNA using Superscript III First-Strand Synthesis kit (ThermoFisher) with primer [Nextera i7 10N] and following the manufacturer’s protocol. cDNA technical replicates were pooled and SPRI Select beads (Life Tech) were used for purification and size selection at a ratio of 0.9X. PCR Library Enrichment was performed using a nested PCR protocol. For the first round of PCR we used Phusion High-Fidelity PCR Master Mix with HF Buffer (NEB) and primers [F trans short and Illumina2.1] with cycling conditions: 98°C for 30s, followed by 15 cycles of 98°C for 10s, 72°C for 15s, followed by 72°C for 5 min. PCR product was purified on a 2% agarose gel as described above. The nested PCR used Phusion High-Fidelity PCR Master Mix with HF Buffer (NEB) and primers [fixed N5xx adapter (Illumina) (unique per each library replicate) and Illumina2.1] with cycling conditions: 98°C for 30s, followed by 5 cycles of 98°C for 10s, 72°C for 15s, followed by 72°C for 5 min. In a side quantitative real-time PCR reaction, 5*μ*L of PCR product, 10X SYBR Green I, and the same primers and master mix were run in conditions: 98°C for 30s, 30 cycles of 98°C for 10s, 63°C for 30s, and 72°C for 60s. To determine the number of PCR cycles needed to reach saturation, we plotted linear Rn versus cycle and determined the cycle number that corresponds to 25% of maximum fluorescent intensity on the side reaction [66]. The PCR product was purified on a 2% agarose gel as described above.

DNA-libraries. We prepared 7 replicates of the DNA library using the PCR protocol as described in [66] except using primers [fixed N5xx adapter (Illumina) (unique per each library replicate) and Nextera i7 10N] and 30ng of input plasmid DNA. PCR product was purified on a 2% agarose gel as described above.

### BiT-BUNDLE-seq

We used BiT-BUNDLE-seq, a new version of the BUNDLE-seq protocol [25]. Input DNA sequences were extracted from the BiT-STARR-seq DNA plasmid library using the same PCR conditions as in preparing the DNA libraries, followed by purification on a 2% agarose gel as described above. We used N-terminal GST-tagged, recombinant human NFKB-p50 subunit from EMD Millipore. The reaction buffer (0.15 M NaCl, 0.5 mM PMSF [Sigma], 1 mM BZA [Sigma], 0.5X TE, and 0.16 *μ*g/*μ*L PGA [Sigma]) was incubated at room temperature for 2 hours in low binding tubes (ThermoFisher). The tubes were cooled for 30 min at 4°C, and then 0.067 *μ*g/*μ*L BSA (Sigma) was added before adding the NFKB-p50 protein. One hundred nanograms of DNA were then added, and the protein and DNA were incubated for 1 h at 4°C. Experiments were performed in triplicates for each NFKB-p50 concentration. The reaction mix was run with 6*μ*L Ficoll (Sigma) in a 7.5% Mini-PROTEAN TGX Precast 10-well Protein Gel (BIORAD) in cold 0.25X TBE buffer for 2 hours at 100V. The gel was stained for 30 min with 3X GelStar (Lonza). Bound and unbound DNA bands were excised under a blue light transilluminator. The DNA was eluted from the gel using the QIAQuick Gel Extraction Kit with a User-Developed Protocol (QIAgen QQ05). The gel slices were incubated in a diffusion buffer (0.5 M ammonium acetate, 10mM magnesium acetate, 1mM EDTA, ph 8.0 [KD Medical]; 0.1% SDS [Sigma]) at 50°C for 30 minutes. The supernatant was then passed through a disposable plastic column containing packed, siliconized glass wool [Supelco] to remove any residual polyacrylamide. Libraries were then quantified and loaded on the NextSeq500 for sequencing.

### Library Sequencing

Pooled RNA and DNA libraries were sequenced on the Illumina Nextseq500 to generate 125 cycles for read 1, 30 cycles for read 2, 8 cycles for the fixed multiplexing index 2 and 10 cycles for index 1 (variable barcode).

### Data Processing

Reads were mapped using the Hisat2 aligner [67], using the 1Kgenomes snp index so as to avoid reference bias. First we removed variants whose UMI was not possible to be present, given the UMI pattern selected. We then ran UMItools [68] using standard flags, as well as a q20 filter. We then ran the deduplicated files through mpileup using a bed file of our full SNP list, the -t DP4, -g, and -d 1000000. DNA reads were processed through a counts filter (on the summed replicates) of more than 7 counts per SNP and at least one count for the reference and alternate alleles in either direction. 50,609 SNPs in the DNA library were used as input to the RNA library. The RNA library was processed following the same procedure as for the DNA library, except that the counts filter required a count of >1 per SNP and at least one count for both reference and alternate alleles. To identify SNPs with allele-specific effects, we applied QuASAR-MPRA [29], where for each SNP the reference and alternate allele counts were compared to the DNA proportion. QuASAR-MPRA results from each replicate were then combined using the fixed effects method, and corrected for multiple tests using BH procedure [69].

### BiT-BUNDLE-seq data analysis

Counts from both the unbound and bound DNA were combined, and a filter was set so that each SNP direction combination had 5 counts for each allele. This combined count was also used to calculate a reference proportion. Each replicate for the bound and unbound libraries were then run through QuASAR-MPRA using the calculated reference proportion. These were then compared using ΔAST [39] to identify ASB in the bound fraction that is differential relative to the unbound fraction. The replicates were combined using Stouffers method [70] to identify ASB for each NFKB-p50 concentration, and combined again to identify the total ASB. The unbound and bound libraries counts were additionally analyzed with DEseq2 [71] to identify over-represented bound enhancer regions (FDR 1% and logFC>1). To better estimate the dispersion parameters, the DESeq2 model was fit on all sequencing data and without merging the replicate libraries:

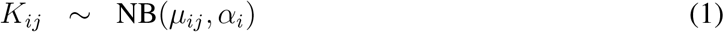

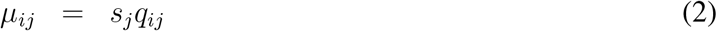

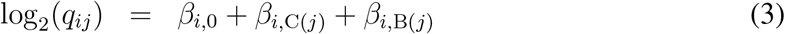

For each enhancer region *i* and sample *j*, the read counts *K_ij_* are modeled using a negative binomial distribution with fitted mean *μ_ij_* and an enhancer region-specific dispersion parameter *α_i_*. The fitted mean is composed of a sample-specific size factor *s_j_* and a parameter *q_ij_* proportional to the expected true concentration of regions for sample *j*. The coefficient *β* _0_ represents the mean effect intercept, *β*_C(*j*)_ represents the lane (NFKB-p50 concentration:replicate) effect, and and *β*_B(*j*)_ represents the Bound/Unbound effect for each NFKB-p50 concentration (High, Medium, and Low).

We then contrasted the bound to the unbound for each concentration (i.e., high concentration bound to high concentration unbound) using the default DEseq2 Wald test for each enhancer region *β*_B(*j*)_ ≠= 0, and a Benjamini-Hochberg (BH) adjusted *p*-value was calculated with automatic independent filtering (DEseq2 default setting).

### GWAS overlap

SNPs nominally significant (p<0.05) for ASB (identified with ΔAST) or ASE (identified with QuASAR-MPRA) that were also annotated as CentiSNP were overlapped with SNPs from the GWAS catalogue (V6) [72], as well as with SNPs fine-mapped with the fgwas software as in [12] with a PPA>0.1.

## Acknowledgement

Funding to support this research was provided by NIH 1R01GM109215-01 (RPR, FL), AHA 14SDG20450118 (FL) and AHA 17PRE33460295 (CK). We would like to thank Wayne State University HPC Grid for computational resources, members of the Luca/Pique group for helpful comments and discussions and Luis Barreiro for making the reQTL data available.

## Supplementary Tables

**Table S1:**
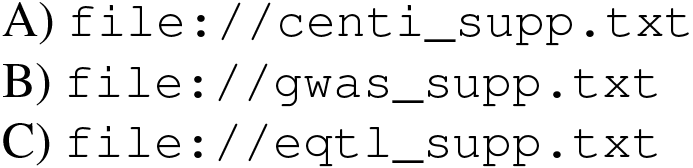
Annotations Used. SNP annotations used for overlap with BiT-BUNDLE-seq and BiT-STARR-seq. First 4 columns are in the same order for each file (chr, pos, pos1, rsID). A) CentiSNPs. Column 5 contains the transcription factor with a CentiSNP at that location. B) SNPs in complex traits. Column 5 contains the GWAS trait associated with the SNP. C) eQTL SNPs. Column 5 contains the information for whether the eQTL was identified in cells infected with L (*Listeria*), S (*Salmonella*), or NI (not infected). Column 6 contains the gene associated with the eQTL. Column 7 contains the beta for the eQTL association. Column 8 contains the *p*-value for the eQTL association.

**Table S2:**
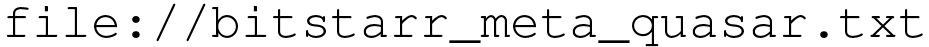
BiT-STARR-seq results. QuASAR-MPRA results for BiT-STARR-seq.

**Table S3:**
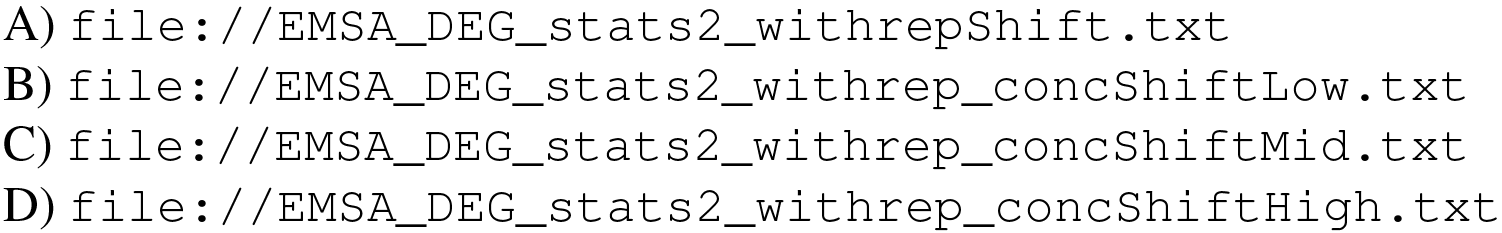
DEseq results. Differentially bound regions for A) Combined concentrations, B) Low concentration, C) Mid concentration, and D) High concentration. Columns are the same for all 4 files (identifier(rsID Direction), adjusted *p*-value, *p*-value, logFC).

**Table S4:**
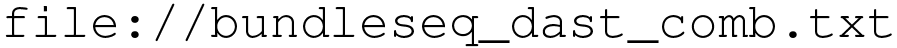
BiT-BUNDLE-seq results. ΔAST results for BiT-BUNDLE-seq. Columns are identifier, z score, *p*-value, adjusted *p*-value, rsID

**Table S5:**
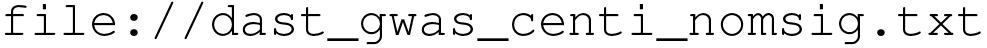
ASB and complex traits. ΔAST results for BiT-BUNDLE-seq. SNPs are nominally significant, associated to a complex trait, and are also CentiSNPs. Columns are rsID, direction, *p*-value, complex trait.

**Table S6:**
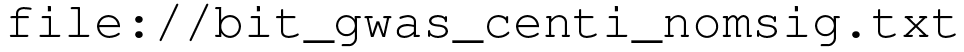
ASE and complex traits. QuASAR-MPRA results for BiT-STARR-seq. SNPs are nominally significant, associated to a complex trait, and are also CentiSNPs. Columns are rsID, direction, *p*-value, complex trait.

**Table S7:** Transcription factors in BiT-STARR-seq. Number of SNPs in motifs matching the top 10 covered transcription factors in BiT-STARR-seq.

| Transcription Factor | Freq |
| --- | --- |
| CTCF | 4911 |
| E2F-1 | 2794 |
| E2F | 4407 |
| ATF | 5567 |
| AML1 | 3794 |
| ATF2:c-Jun | 3651 |
| CREB | 12955 |
| AP1 | 2673 |
| ARG RI | 3445 |
| STF1 | 3561 |

**Table S8:**
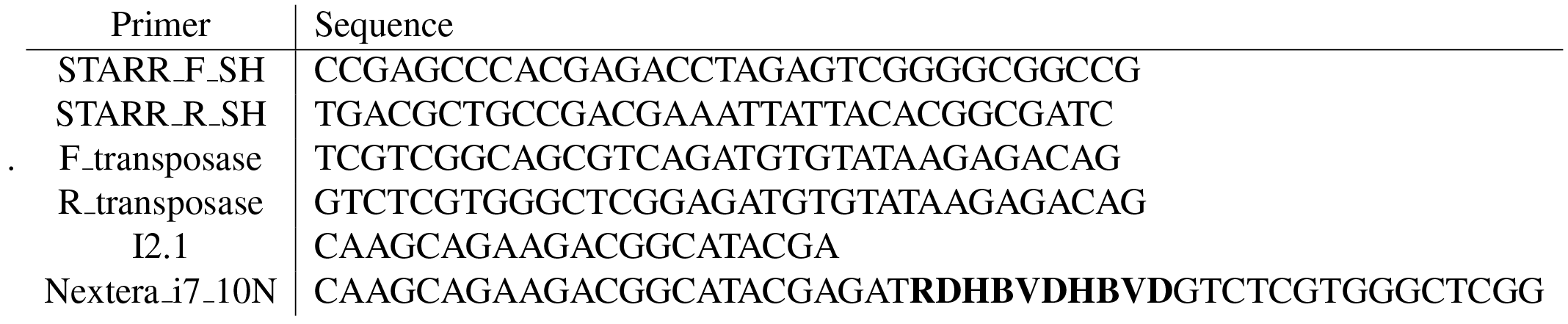
Primers used in BiT-STARR-seq.

## Supplemental Figures

**Figure S1:**
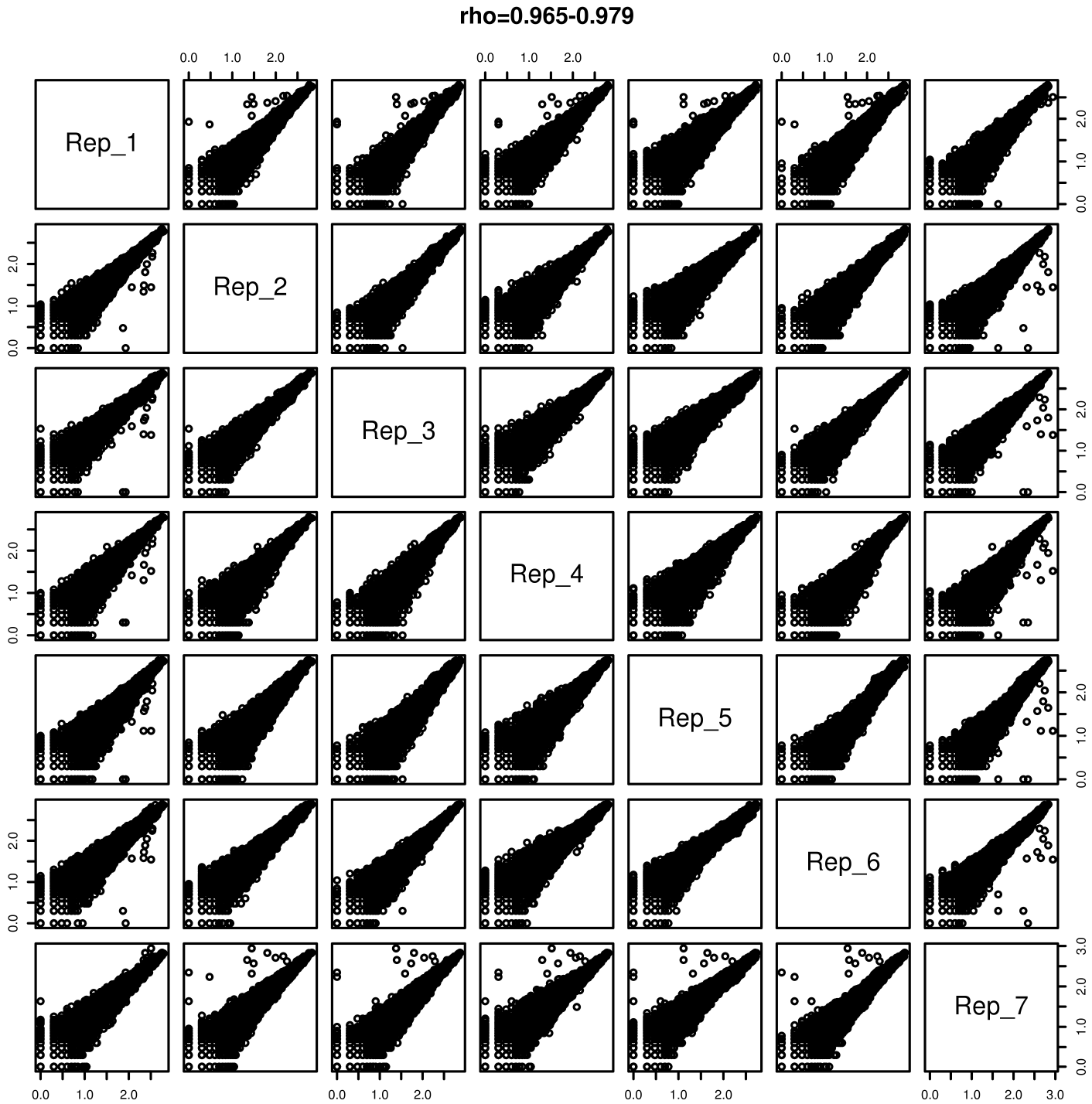
Correlation of DNA libraries. Scatterplot of filtered DNA library counts for each replicate plotted against all other replicates. Spearman *rho* correlation range is stated at the top.

**Figure S2:**
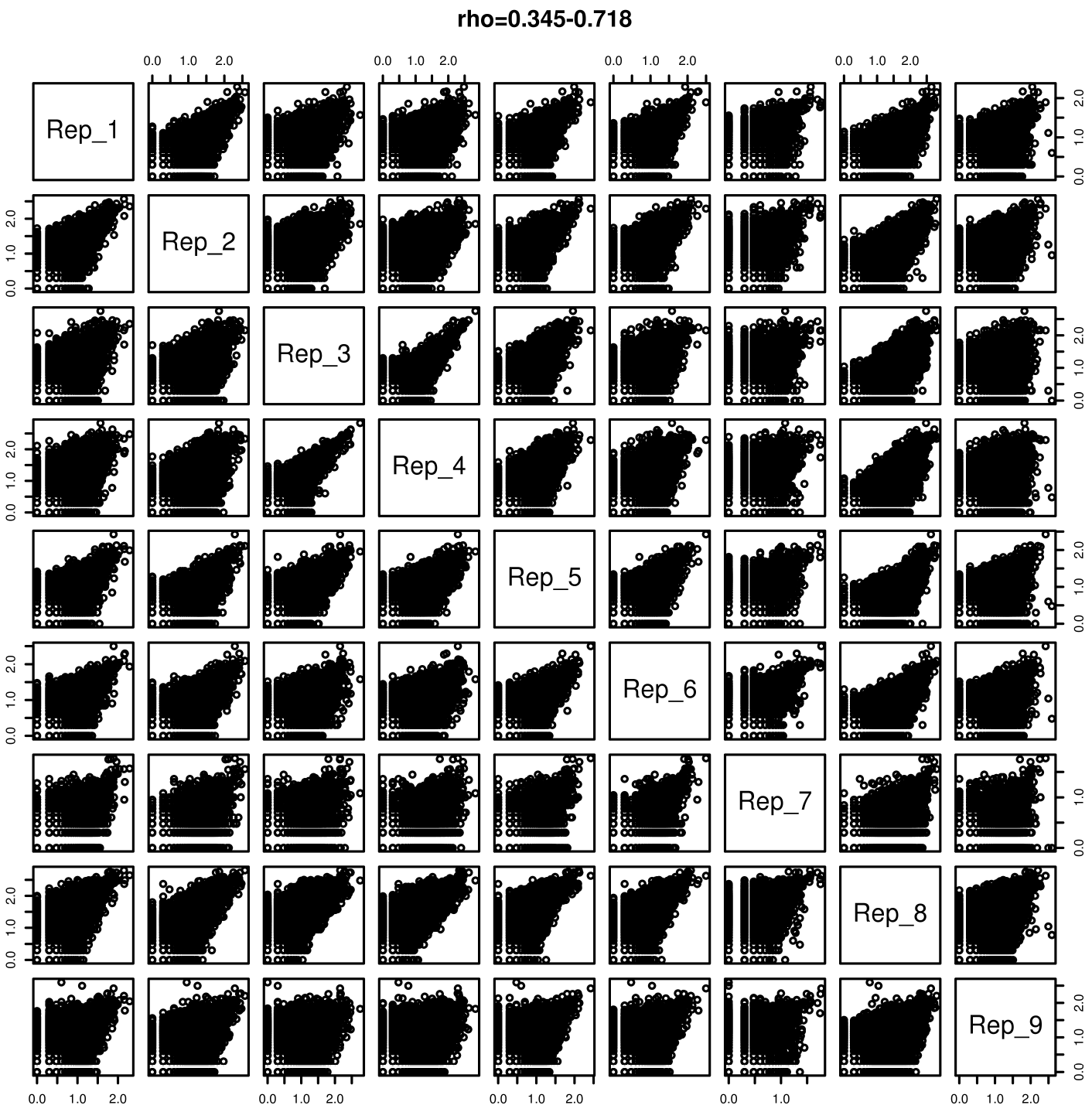
Correlation of RNA libraries. Scatterplot of filtered RNA library counts for each replicate plotted against all other replicates. Spearman *rho* correlation range is stated at the top.

**Figure S3:**
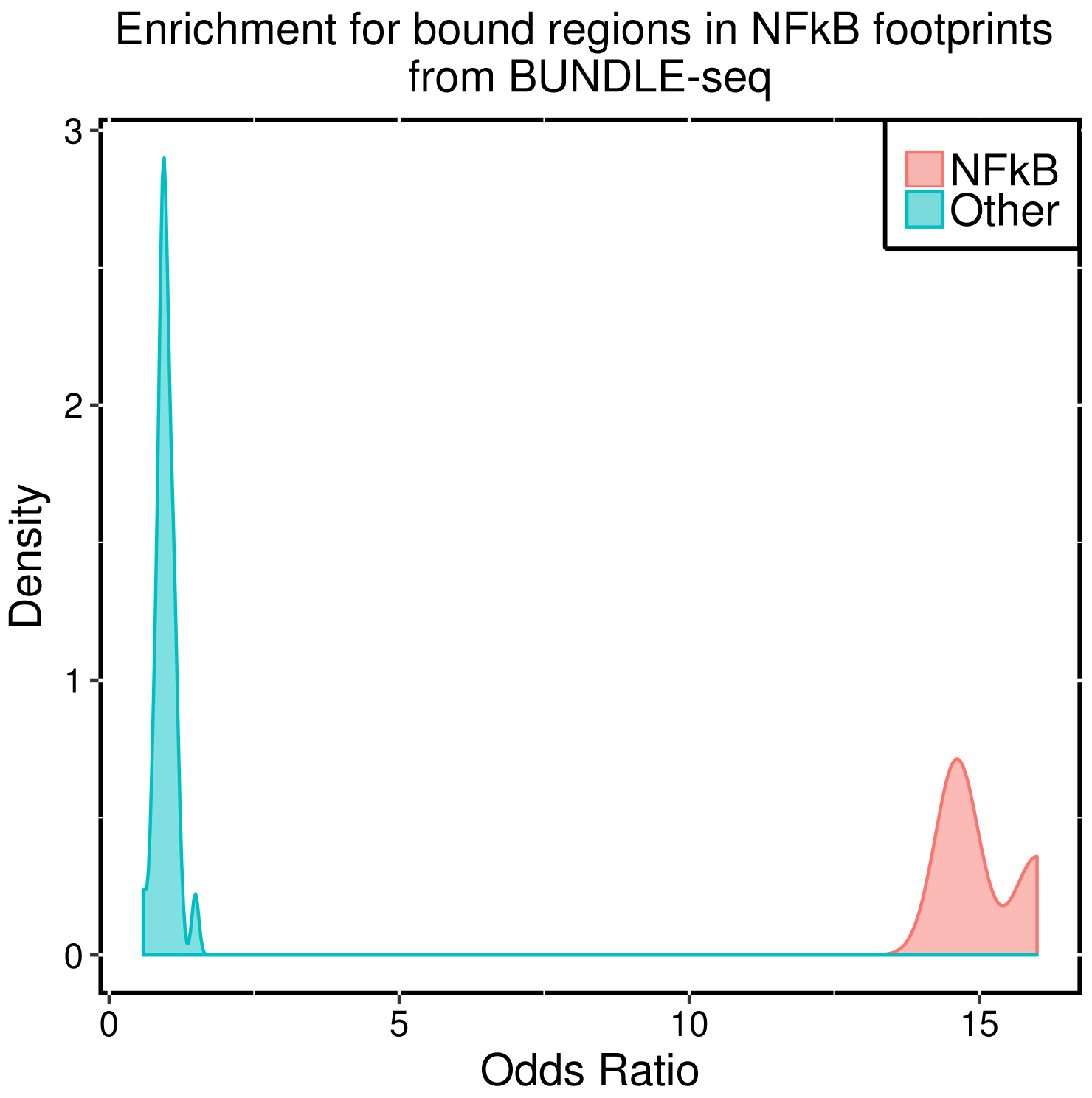
Enrichment of NFKB footprints in BiT-BUNDLE-seq bound regions. Fishers exact test was performed to identify enrichment (x axis is the OR) for significant differentially bound regions (logFC>1 and FDR<1%). In red are the regions containing a SNP in a NFKB footprint, in blue the regions containing a SNP in footprints for other transcription factors.

